# A quantitative comparison of two methods for higher-order EEG microstate syntax analysis

**DOI:** 10.1101/2025.06.11.658518

**Authors:** Frederic von Wegner, Gesine Hermann, Inken Tödt, Inga Karin Todtenhaupt, Helmut Laufs

## Abstract

Entropy rate (ER) and sample entropy (SE) are two metrics that have been used to quantify the syntactic complexity of electroencephalography (EEG) microstate sequences. We here present a theoretical and numerical comparison of these two metrics and apply them to a resting-state EEG dataset from individuals with Alzheimer’s disease (AD) and a control group.

We first derive theoretical entropy rate and sample entropy estimates for first-order discrete Markov processes, providing a null hypothesis for statistical testing of higher-order syntax properties. Under the first-order syntax null hypothesis, we find a close mathematical relationship between both metrics that can be expressed by the microstate transition probability matrix. An inequality is derived that shows entropy rate to be an upper bound to sample entropy under the Markov approximation.

We quantify accuracy and precision of the theoretical ER and SE estimates on EEG microstate sequences from the healthy control group. We then show that ER and SE identify significant higher-order syntax properties in microstate sequences from the control and AD groups. Group comparison demonstrates that continuous microstate sequences from the AD group have lower entropy values (ER, SE), whereas jump sequences from the AD group have higher entropy values compared to control. Finally, we introduce a new syntax metric that normalizes ER and SE values with respect to their first-order syntax levels, to assess differences that only depend on syntax order. This metric revealed no differences between control and AD groups for either continuous or jump microstate sequences.

This study provides further insights into higher-order microstate syntax and how it can be quantified with respect to the underlying first-order syntax. Similarities and differences between ER and SE as syntax metrics are highlighted and exemplified on experimental data. Our results show that (i) EEG microstate sequences from control and AD subjects show higher-order syntax properties across the tested syntax levels, (ii) continuous and jump sequences from control and AD groups are syntactically different, and (iii) differences between the control and AD groups disappear when higher-order syntax properties are normalized to the group-specific Markov level.

## 1 Introduction

EEG microstate analysis is a versatile tool to quantify spontaneous and evoked EEG dynamics (Michel and Koenig, 2018). Continuous-time, multi-channel EEG data is compressed into a sequence of labels taken from a small alphabet, where each label refers to a cluster of topographically similar EEG data vectors. The interpretation is that each cluster corresponds to a functional brain network (Michel and Koenig, 2018) and it has been suggested that changes in topography and dynamics correlate with cognition (Britz et al, 2010; Custo et al, 2017; Seitzman et al, 2017) and a variety of brain pathologies (Smailovic et al, 2019). Historically, microstate sequences have most commonly be quantified by the three parameters, duration, occurrence and coverage. Lehmann et al (2005) introduced a syntax analysis based on single time step transition dynamics, and the same study also addressed differences in four-step transitions between individuals with schizophrenia and healthy controls. The single time step, or first-order, syntax concept has since been used to characterize a wide range of experimental conditions and brain sates, among them cognitive tasks (Seitzman et al, 2017; Antonova et al, 2022), vigilance and sleep (Brodbeck et al, 2012; An et al, 2024), schizophrenia (Lehmann et al, 2005; Tomescu et al, 2015), and Alzheimer’s disease (Nishida et al, 2013; Musaeus et al, 2019, 2020; Tait et al, 2020).

Although first-order syntax analysis has generated many new insights, it sometimes does not reveal significant changes despite measurable neurological or cognitive symptoms (Nishida et al, 2013; Musaeus et al, 2019; Tait et al, 2020), and this has generated a growing interest in higher-order syntax methods (von Wegner et al, 2017; Tait et al, 2020; Murphy et al, 2020; Wiemers et al, 2023; Hermann et al, 2024; Artoni et al, 2023; von Wegner et al, 2023; Haydock et al, 2025). The common target of these methods are multi-time-step microstate transition dynamics, which are commonly described by multivariate discrete distributions over the possible microstate words of a given length (von Wegner et al, 2025; Haydock et al, 2025). To avoid a brute-force enumeration of these extensive microstate word dictionaries, different functionals are applied to these distributions to obtain a scalar quantity that is easier to interpret. Following this definition, higher-order syntax comprises approaches such as (finite) entropy rate (von Wegner et al, 2018, 2023; Hermann et al, 2024), excess entropy (von Wegner et al, 2023), active information storage (Hermann et al, 2024), Microsynt (Artoni et al, 2023), sample entropy (Murphy et al, 2020; Teng et al, 2024; Wu et al, 2025), and epsilon machines (Haydock et al, 2025).

The aim of this article is to give a quantitative analysis and comparison of two of these metrics, entropy rate and sample entropy. This study is motivated by the fact that, while we have used entropy rate to quantify microstate dynamics in the past (von Wegner et al, 2018, 2023; Hermann et al, 2024), others have used sample entropy (Murphy et al, 2020; Teng et al, 2024; Wu et al, 2025), and both are commonly interpreted in a similar way, namely as a measure of sequence syntax or complexity. Addressing a similar question, we have demonstrated an equivalence between entropy rate and Lempel-Ziv complexity in a recent study (von Wegner et al, 2023), considerably facilitating the interpretation and comparison of different studies. The question of this study is whether entropy rate and sample entropy yield equivalent results, a finding that would further facilitate the comparison of existing studies and the design of future studies. Equivalence cannot be assumed on theoretical grounds, as these metrics describe different dynamical concepts. Furthermore, these quantities have been established in the field of real-valued, continuous dynamical and stochastic systems (Gaspard and Wang, 1993), whereas EEG microstate analysis concerns discrete-valued time series. The discrete nature of microstate sequences offers an opportunity to arrive at approximate or even exact results for ER and SE, and thus a better understanding of these syntax metrics. This article has two aims. The first aim is to establish a quantitative relationship between ER and SE under first-order Markov assumptions. A quantitative understanding of first-order syntax will be useful in deriving null hypotheses for higher-order syntax testing. The second aim is to test the theoretically expected results and to search for higher-order microstate syntax on a publicly available EEG dataset of resting-state recordings from healthy control subjects (cognitively normal, CN) and individuals with Alzheimer’s disease (AD) (Miltiadous et al, 2023).

The article is structured as follows. We briefly review the definitions of entropy rate and sample entropy and their simplified versions in the case of discrete signals and then derive ER and SE estimates under a first-order Markov assumption. We then test the accuracy of these estimates on empirical microstate sequences from healthy subjects. Higher-order syntax is statistically assessed in the CN and AD groups separately, and finally the two groups are compared directly. All tests are applied to continuous-time microstate sequences and jump sequences, as the known non-Markovian features of microstate sequences prevent us from predicting jump sequence properties from continuous sequences (von Wegner et al, 2017).

## 2 Background

### 2.1 Microstate syntax

For quantitative analysis, microstate sequences are interpreted as stochastic processes, i.e. as sequences of random variables *X*_*t*_ with discrete time index *t*. The syntactic structure and complexity of these sequences can be quantified in terms of temporal correlations between different time points, and can be described systematically by discrete Markov chain models of a given order. At the same time, these models provide null hypotheses for statistical testing.

We write the set of *K* microstate labels as l*K*J = {0, 1, …, *K* − 1} and, where useful, we implicitly convert between integers and the microstate labels A-D as used in the literature (Michel and Koenig, 2018). Array indices can thus be identified with microstate classes directly. With this indexing, the microstate distribution **p** = (*p*_0_, …, *p*_*K*−1_) is an array where *p*_*i*_ is the probability of microstate class *i* in the sequence *X*_*t*_, i.e. *p*_*i*_ = Pr (*X*_*t*_ = *i*). The conditional transition matrix **T** is a two-dimensional array that contains the conditional transition probabilities

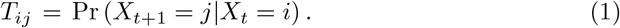

We are also going to use the closely related matrix **P** of joint (instead of conditional) transition probabilities

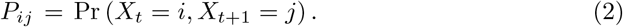

Both matrices are equally useful to desribe first-order properties and they can be converted into each other by

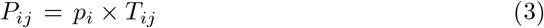

but readers should be aware that both **T** and **P** have been called transition matrix in the microstate literature (von Wegner et al, 2017; Hermann et al, 2024; Al Zoubi et al, 2019; Antonova et al, 2022; Tomescu et al, 2015).

The most common choice of microstate syntax analysis is a description of first-order properties (Lehmann et al, 2005) in terms of the microstate distribution **p** and the transition matrix **T** (von Wegner et al, 2025).

Higher-order microstate syntax describes statistical dependencies of *X*_*t*+1_ on longer microstate histories of length *k* (Hermann et al, 2024), also called ‘words’ in Artoni et al (2023); von Wegner et al (2025) or ‘n-grams’ in Haydock et al (2025). A k-history (k-word) can be written as

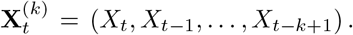

For a Markov process of order *m*, the prediction of the next microstate *X*_*t*+1_ depends only on the last *m* states (Billingsley, 1961):

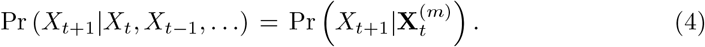

A zero-order Markov process (*m* = 0) is defined by its distribution **p** only and has no temporal correlations, and a first-order Markov chain is fully characterized by its transition matrix **T**. We therefore define higher-order syntax as any microstate sequence property described in terms of a conditional distribution Pr 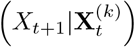 for *k >* 1 that cannot be predicted by the first-order Markov properties **p** (microstate distribution) and **T** (transition matrix) (von Wegner et al, 2025).

### 2.2 Entropy

To obtain a more easily interpretable scalar quantity, different functionals can be applied to the higher-order transition matrix 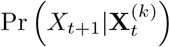. Two particular choices of such functionals result in the quantities entropy rate (ER) and sample entropy (SE). ER and SE have been applied to continuous dynamical systems, physiological time series (Eckmann and Ruelle, 1985; Gaspard and Wang, 1993; Richman and Moorman, 2000; Lake et al, 2002), and more recently to EEG microstate sequences (von Wegner et al, 2023; Hermann et al, 2024; Murphy et al, 2020; Teng et al, 2024; Wu et al, 2025).

#### 2.2.1 Entropy rate

Entropy rate measures how much uncertainty (randomness) a system produces with each observed sample. This can be reformulated as the predictability of the future state *X*_*t*+1_, given knowledge of the system’s past states (*X*_*t*_, *X*_*t*−1_, …). To allow computability and to test the dependency on finite history lengths, the set of past states is truncated to a finite number of *k* time steps. A finite estimate of entropy rate for word (history) length *k* ≥ 0 is therefore given by

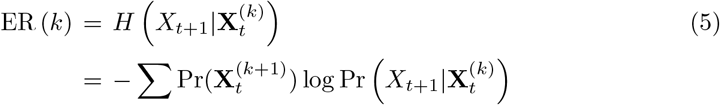

where the second line applies the definition of conditional entropy (Cover and Thomas, 2005) and the sum ranges over all possible microstate words 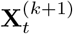.

Since ER (*k*) is computed from the conditional distribution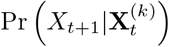, it is a higher-order syntax measure according to our definition (von Wegner et al, 2025). As a side note, our notation ER (*k*) should not be confused with Eckmann-Ruelle entropy, sometimes abbreviated E-R entropy (Costa et al, 2005; Delgado-Bonal and Marshak, 2019).

#### 2.2.2 Sample entropy

Sample entropy (SE) (Richman and Moorman, 2000) was introduced as an alternative to approximate entropy (Pincus et al, 1991; Pincus and Goldberger, 1994), and both are different from entropy rate (Pincus and Goldberger, 1994; Costa et al, 2005). SE is also based on the higher-order transition matrix 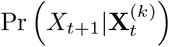, but different from ER, it quantifies the return of microstate words of length *k* after an arbitrary time interval. The k-th sample entropy coefficient SE (*k*) is related to the probability that, given the recurrence of a microstate word of length *k* at time points *t* and *s*, the recurrence extends to the corresponding *k* + 1 word. Practically, given a template word 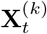, the remaining microstate sequence is scanned for another recurrence of the same word. If such a recurrence is found at time *s*, i.e. 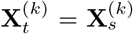, it is tested whether *X*_*t*+1_ = *X*_*s*+1_. If this is true, the recurrence remains valid for the longer word 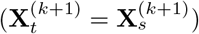 and indicates a more regular sequence, i.e. more syntactic structure. The algorithm estimates the conditional probability 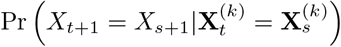 which is turned into an entropy-like quantity by taking the negative logarithm:

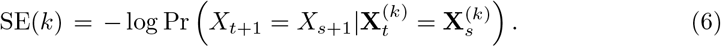

Higher SE values reflect less syntactic structure and more randomness. Sample entropy is related to the Rényi entropy of order two, also known as collision entropy (Costa et al, 2005). The concept is illustrated for the microstate context in Figure 1, using a two-dimensional (2D) projection for visualization. The original EEG data vectors were clustered with the modified K-means algorithm (K=4) (Pascual-Marqui et al, 1995). 2D-coordinates of EEG data vectors at global field power (GFP) peaks were obtained from principal component analysis (PCA) and are shown as coloured dots in Figure 1. The four colours indicate the cluster assignment according to the K-means output (microstate class A: black, B: blue, C: red, D: yellow). The solid black line is the 2D projection of 200 ms of continuous EEG data. Individual samples are shown as open circles and the arrow of time is indicated by the two blue solid arrows. The associated microstate sequence can be inferred from the microstate ‘sector’ on which the trajectory falls. This corresponds to the mathematical back-fitting process during which microstate templates are mapped into the EEG dataset. Two microstate words 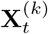 and 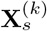 are highlighted with filled circles. They can be parametrized as 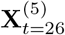 and 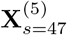 within the short example sequence and refer to the same microstate word BBBBA, so 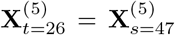. For convenience, the microstate word is written with time running from left to right. Entropy rate measures how much information the preceding four B’s encode about the subsequent state A. The two words 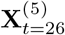 and 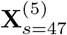 enter the entropy rate calculation independently. If one of the words was deleted, the other word would still have the same contribution to the ER calculation, but both words are needed to calculate SE. In this sense, ER (*k*) measures the short-term predictability of the sequence, based on the last *k* samples. Sample entropy, in contrast, relies on the recurrence of a given microstate word anywhere along the sequence. The re-entry of the EEG trajectory into the same small region of the 2D state space explains the term ‘collision entropy’. The original SE algorithm for continuous systems (Richman and Moorman, 2000) relies on a finite radius that defines the size of the neighbourhood

**Fig. 1.**
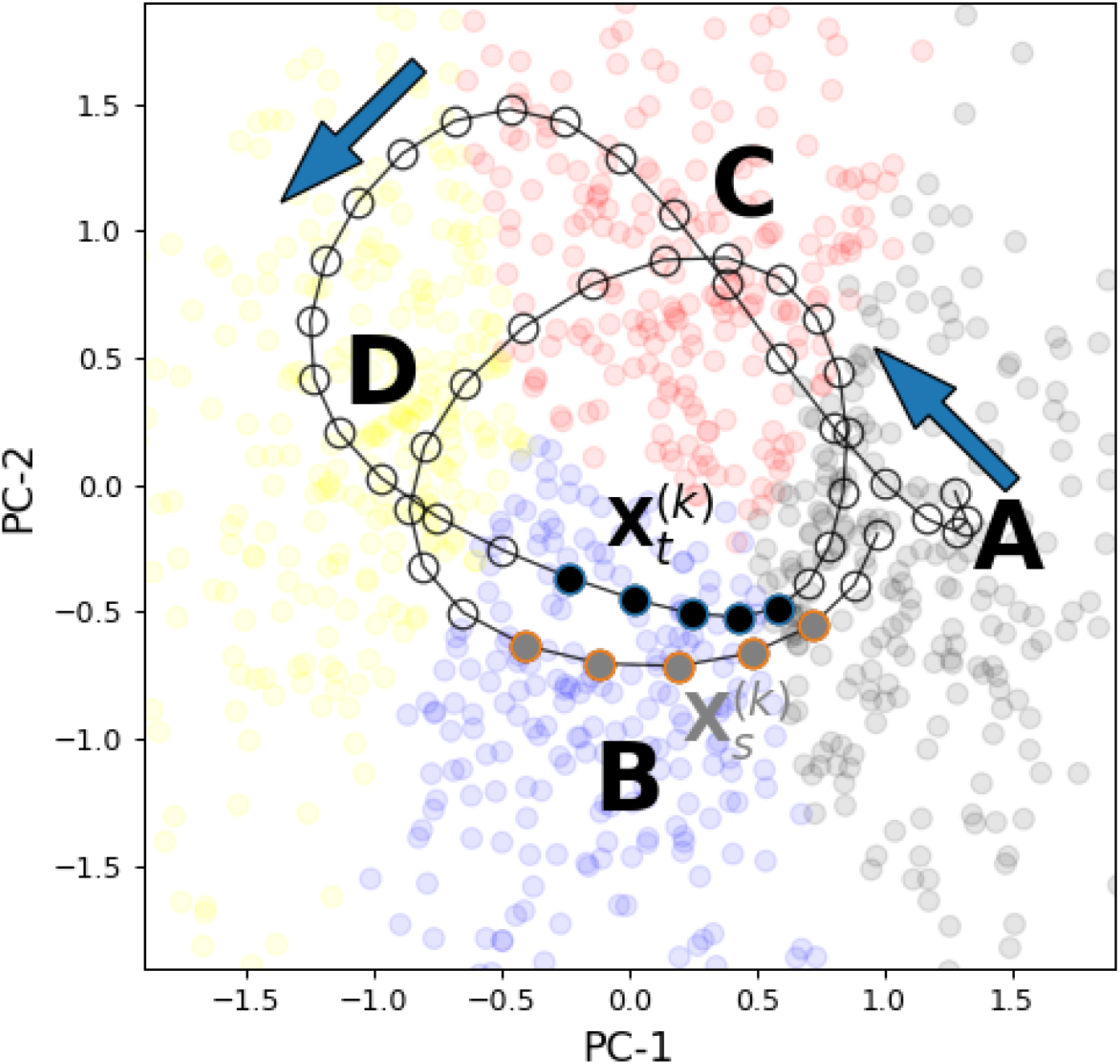
Higher-order microstate sequence syntax. An EEG segment and its corresponding microstate sequence are illustrated as follows: coloured dots in the background show a 2D-PCA projection of the EEG data vectors at GFP peaks from which microstate maps were obtained through modified K-means clustering. Colour indicates the microstate class to which each GFP peak was assigned (microstate class A: black, B: blue, C: red, D: yellow). The 2D-PCA projection of 200 milliseconds of continuous EEG trajectory is overlaid (empty dots and solid line, arrows indicate time). A quasi-periodic structure of the trajectory is observed. The shown EEG segment translates into the continuous microstate sequence A^6^C^6^D^9^B^5^A^5^C^7^D^4^B^5^A^3^ and the microstate jump sequence ACDBACDBA. The two microstate words 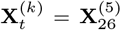 and 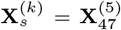 are identical (BBBBA, time running from left to right) and are interpreted as a ‘collision’ of two trajectory segments. The high-lighted words 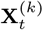 and 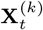 end with the first and second BA transition of the sequence above, respectively.

which qualifies as a collision. This is usually expressed in terms of the continuous signal’s standard deviation. For discrete microstate sequences, although this parameter is sometimes mentioned in the microstate literature (Teng et al, 2024; Wu et al, 2025), it does not have to be provided, and if provided, should not affect the output of the algorithm. For microstate sequences, a collision is defined as exact equality between two microstate words.

#### 2.2.3 Entropy rate and sample entropy estimates for Markov processes

This section contains new results regarding the theoretically expected ER and SE values for first-order Markov processes. Due to its theoretical nature and its close link to the preceding sections, these results are summarized here, instead of the Results section where analyses of actual EEG data are presented.

Under the assumption that *X*_*t*_ is a first-order Markov process, an ER estimate for finite history length *k* is obtained as:

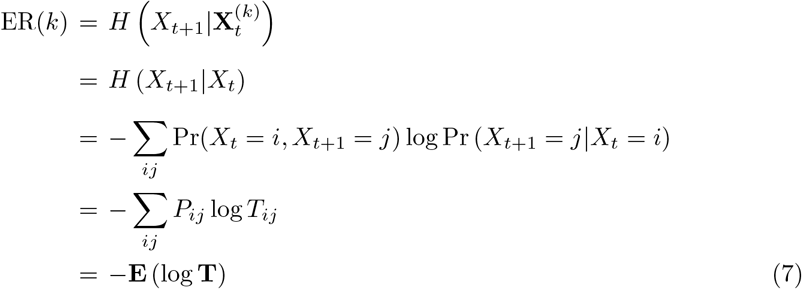

where we have used Equation 5 (line 1), the first-order Markov property (line 2), the conditional entropy definition (line 3) (Cover and Thomas, 2005), the expressions for joint (**P**) and conditional (**T**) first-order transition matrices (line 4), and **E**(·) as the expected value of a random variable with respect to the distribution **P** (line 5).

For the k-th sample entropy coefficient, we use an approximation:

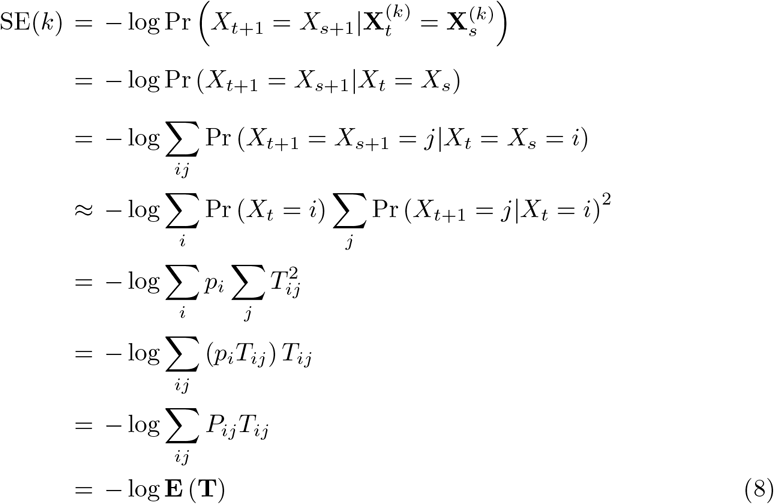

where we have used the SE definition (Equation 6, line 1), truncation as justified by the Markov assumption (line 2), an independence assumption further discussed below (line 4), the definitions of microstate distribution (**p**), joint (**P**) and conditional (**T**) transition matrices (lines 5-7), and the expectation **E**(·) is again computed with respect to **P**. A key step of this approach is the approximation in line 4, that collisions are independent for all recurrence intervals |*t* − *s*|, which seems justified when the recording is much longer than the sequence autocorrelation time. Due to independence, the conditional probability of two *i* → *j* transitions, one at time *t* and the other at time *s*, can be expressed as 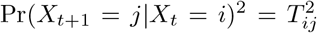. This probability is conditional on the preceding microstate words 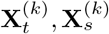 terminating in *X* = *X* = *i*, hence the weighting of 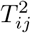 by *p*_*i*_. Independence along the time axis is often assumed for time lags beyond the first minimum of the autocorrelation or autoinformation function (Fraser and Swinney, 1986), which is located near 25 ms for resting-state EEG microstate sequences (von Wegner et al, 2017). The accuracy of the approximation is tested numerically against Markov surrogate data in the Results section. Given the concavity of the logarithm function, the two results can be summarized as an inequality:

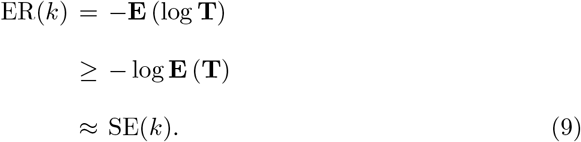

Equation 9 shows that entropy rate and sample entropy for finite word lengths are closely related quantities in the first-order Markov case. Their difference lies in the order in which expectation and logarithm operations are applied to the entries of the (conditional) transition matrix **T**. The finite entropy rate estimate ER(*k*) provides an upper bound for the corresponding sample entropy coefficient SE(*k*).

In the following, all logarithms will be taken to the base 2, i.e. log = log_2_.

## 3 Data and Methods

### 3.1 EEG

We analyzed eyes-closed resting-state EEG recordings from a public EEG database described in Miltiadous et al (2023). The dataset contains healthy control subjects and two patient groups, one group diagnosed with Alzheimer’s disease, the other with fronto-temporal dementia. We here describe properties of the control group (CN) and the Alzheimer’s disease group (AD). The published pre-processed data is 19-channel EEG data sampled at 500 Hz and band-pass filtered in the 0.5-45 Hz range.

The pre-processed data has undergone artefact correction with the Artifact Subspace Reconstruction (ASR) method (Chang et al, 2018) and subsequent independent component analysis (ICA). The detailed procedures are described in Miltiadous et al (2023).

### 3.2 Microstate analysis

The pre-processed published data contains discontinuities where artefact-rich intervals have been excluded. We used the annotated discontinuities to split each recording into contiguous segments and only considered the longest segment of each subject for analysis. Since higher-order syntax analysis requires a large number of samples, we only included microstate sequences with at least 10^5^ samples, and truncated longer recordings at 10^5^ samples (200 seconds) to ensure uniformity across subjects. This was fulfilled for n=24/29 CN subjects and n=25/36 AD subjects.

We used MNE (Gramfort, 2013) and the Pycrostates package (Férat et al, 2022) for microstate analysis. Each EEG data set was band-pass filtered to the 1-30 Hz frequency band (zero-phase, non-causal digital filter, -6dB cutoff frequencies 0.5 Hz and 33.75 Hz) and an average reference was computed. We used modified K-means clustering of EEG data vectors at temporal maxima of the global field power time series to obtain K=4 template maps per subject, followed by group-level modified K-means clustering of the subject template maps (Pascual-Marqui et al, 1995). Group-level maps used in this article were obtained from EEG recordings across all three groups. The group-level maps were labelled manually according to standard conventions (Michel and Koenig, 2018). Clustering quality was similar for different choices of K. We selected K=4 to simplify the presentation.

Microstate sequences were obtained by competitive back-fitting of the group-level maps at each time point, followed by smoothing with regularization parameter *λ* = 5 and window half-size of *b* = 3 samples, resulting in a smoothing window size of 14 milliseconds (Pascual-Marqui et al, 1995). The first and last segment of each microstate sequence were excluded. In analogy to continuous-time Markov chain theory, these sequences will be referred to as continuous microstate sequences. Removing adjacent duplicate labels yields the corresponding jump sequence (von Wegner et al, 2023).

### 3.3 A normalized syntax measure

We here introduce a normalized syntax measure for continuous and jump sequences. The aim of this new metric is to analyze the shape of ER and SE curves, independent of their ‘offset’, i.e. their absolute entropy levels at *k* = 0 and *k* = 1, reflecting zero- and first-order Markov properties, respectively. In contrast to continuous microstate sequences, jump sequences cannot have zero-order syntax (von Wegner et al, 2025). We therefore include zero and first-order coefficients into the normalized syntax metrics for continuous sequences and first-order coefficients for jump sequence syntax metrics. The following definitions can be applied to both ER and SE coefficients. All terms *h* below can therefore represent either ER or SE values.

Thus, for continuous sequences with theoretical entropy values of *h*_0_ and *h*_1_ under zero- and first-order Markov assumptions, respectively, and empirical entropy value *h* for the real EEG microstate sequence, respectively, we define the normalized syntax metric as the function:

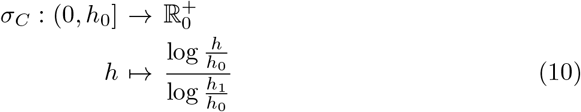

For jump sequences, that do not have a zero-order Markov model, we define:

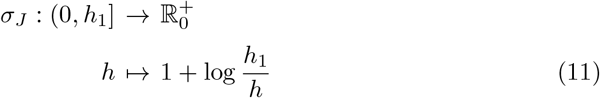

The two metrics fulfill:

1. *σ*_*C*_ (*h*) = 0 if the (continuous) sequence behaves like a zero-order Markov model (maximum entropy *h* = *h*_0_).
2. *σ*_*C,J*_ (*h*) = 1 if the (continuous or jump) sequence behaves like a first-order Markov model (*h* = *h*_1_).
3. *σ*_*C,J*_ (*h*) *>* 1 if the (continuous or jump) sequence has more syntactic structure (is more predictable) than a first-order Markov model (*h < h*_1_).
4. *σ*_*C,J*_ are monotonically increasing.

### 3.5 Surrogate data

We generated zero-order and first-order Markov chain surrogate sequences using a previously published method (von Wegner and Laufs, 2018; von Wegner et al, 2025). Surrogate sequences have the same number of samples as the original sequences. Zero-order surrogates have the same microstate distribution **p** as the original sequence *X*_*t*_ and transition matrix coefficients *T*_*ij*_ = *p*_*j*_. First-order surrogates have the same microstate distribution **p** and transition matrix **T** as the actual EEG microstate sequence.

### 3.5 Statistics

When comparing sets of ER or SE values for a given syntax order against Markov surrogates within a group (CN or AD), we applied the Wilcoxon test for dependent samples. When comparing ER or SE values for a given syntax order between CN and AD groups, we applied the Mann-Whitney U-test for two independent samples and unequal sample sizes. The confidence level was set to *α* = 0.05.

## 4 Results

### 4.1 Theoretical and numerical estimates for Markov surrogates

We tested the accuracy of the theoretical estimates given in Equation 7, Equation 8 and Equation 9 for word lengths *k* = 1 − 6 numerically, using Markov surrogate data. The results are shown in Figure 2. Surrogates had the same number of samples as real microstate sequences, i.e. 10^5^ samples for continuous sequences, and an average of 8564 samples (range: 7198-9766) for jump sequences. In Figure 2, theoretical ER (SE) coefficients calculated from Equation 7 (Equation 8) are shown as dotted lines (label ‘theo.’). The values are defined for integer values of *k* only, but are shown as continuous lines for the purpose of illustration. Coefficients calculated from surrogates (label ‘surr.’) with either zero-order (MC-0, red) or first-order (MC-1, blue) syntax are shown as open circles with error bars (mean, standard deviation).

**Fig. 2.**
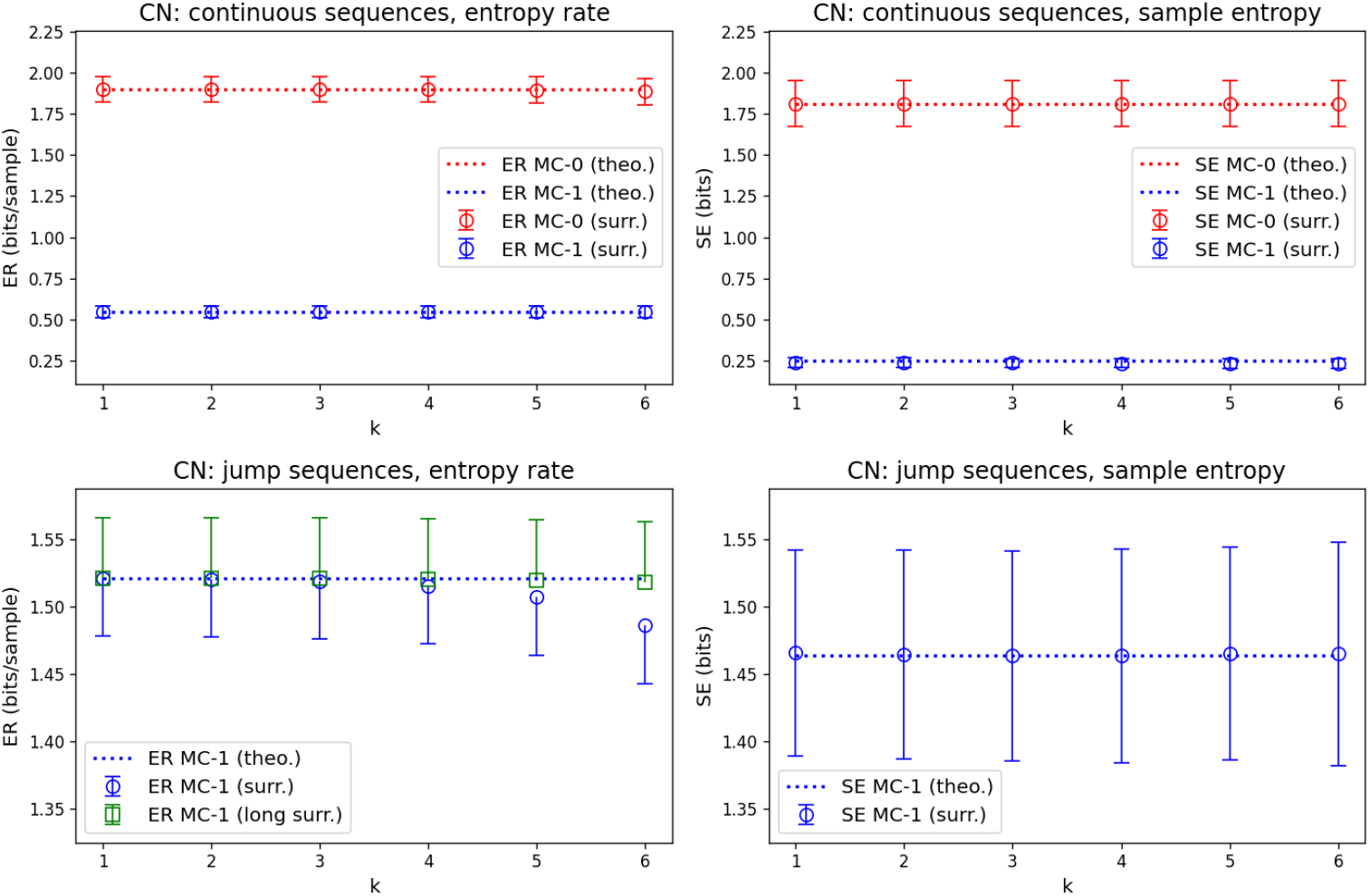
Accuracy of theoretical approximations for entropy rate (Equation 7) and sample entropy (Equation 8) under Markov assumptions. For each EEG microstate sequence from the control group (CN, n=24 subjects), theoretical ER and SE values under zero-order (red dotted, label: MC-0 (theo.)) and first-order (blue dotted, label: MC-1 (theo.)) Markov assumptions were averaged across subjects and compared to the corresponding Markov surrogate sequences. Continuous sequences had 10^5^ samples, jump sequences had 8564 samples on average (range: 7198-9766 samples). Continuous sequences have zero-order (red, label: MC-0 (surr.)) and first-order (blue, label: MC-1 (surr.)) Markov surrogates, jump sequences only have first-order surrogates (blue, label: MC-1 (surr.)). The ER bias observed for shorter jump sequence surrogates (blue circles, label: ER MC-1 (surr.)) was removed when a set of longer surrogate sequences (10^5^ samples, green squares, label: ER MC-1 (long surr.)) was analyzed.

For continuous sequences, ER and SE estimates show a high degree of concordance between theoretical estimates and numerical Markov chain surrogate sequences. Fulfillment of the inequality ER (*k*) ≥ SE (*k*) (Equation 9) can also be observed.

For the entropy rate of jump sequences, Markov surrogates showed a good agreement for short words (*k* = 1 − 4) but showed a bias towards lower ER values for higher-order syntactic properties (*k* = 5, 6). We hypothesized that the bias was caused by an insufficient number of samples in the jump sequences. This was confirmed by analyzing another set of Markov surrogates for jump sequences that had 10^5^ samples per sequence. These longer surrogates showed a good agreement with theoretical ER estimates (green squares and error bars).

### 4.2 Control vs. Alzheimer’s disease group

This section contains three analyses: (i) higher-order syntax analysis for the CN group; (ii) higher-order syntax analysis for the AD group; (iii) a direct comparison between CN and AD groups.

Figure 3 shows the syntax analysis for the CN group (black lines and open triangles) as compared to first-order Markov surrogates (blue lines and open circles). For each of the n_CN_=24 control subjects, SE and ER coefficients were computed for each real microstate sequence and a surrogate sequence of equal length, and the results are illustrated as means (open symbols) and standard deviation (half error bars).

**Fig. 3.**
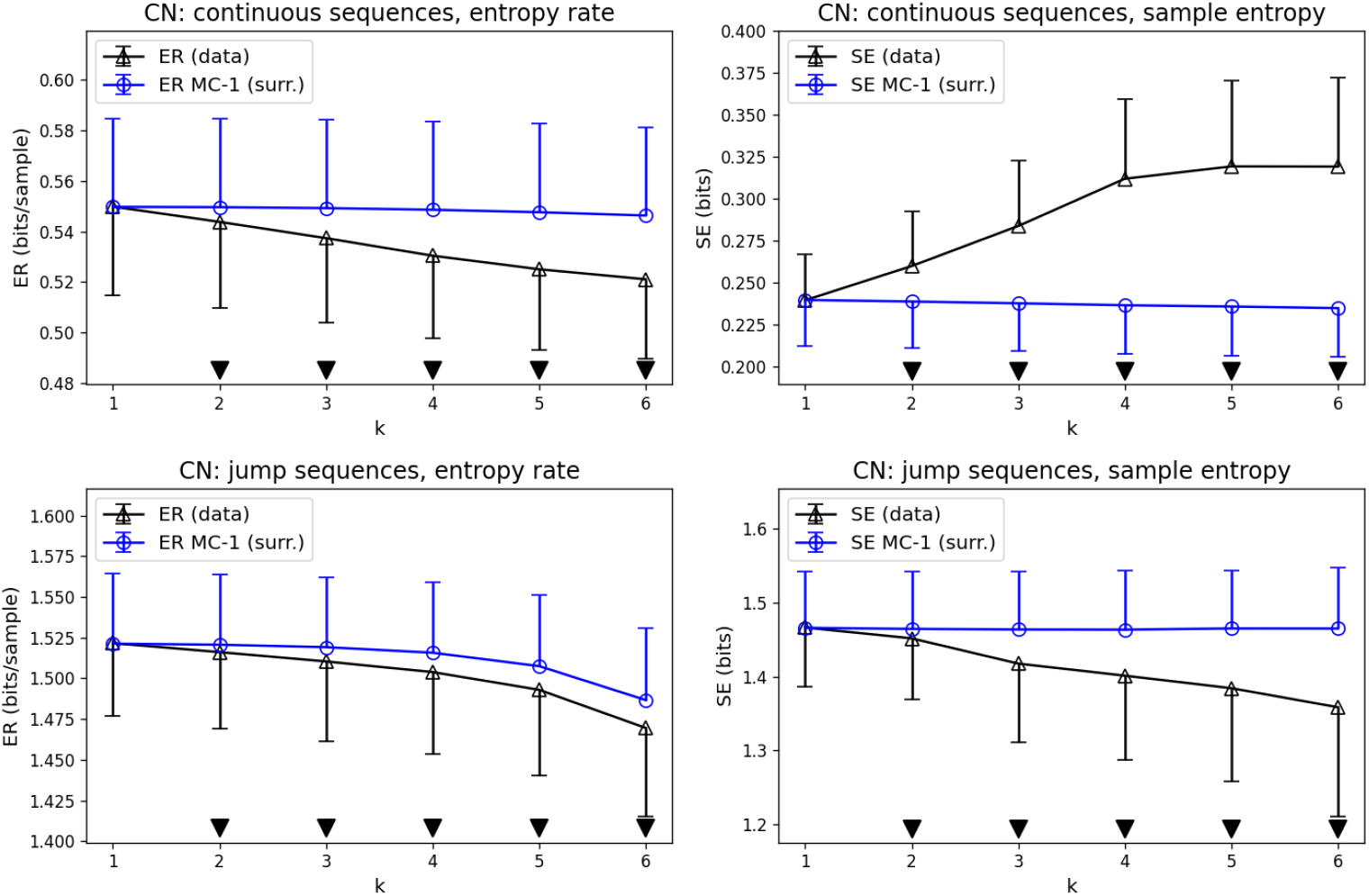
Entropy rate (ER) and sample entropy (SE) compared for real microstate sequences from the CN group (black, label: data) and their first-order Markov surrogates (blue, label: MC-1 (surr.)). Statistically significant differences (p *<*0.05) between real microstate sequences and first-order Markov surrogates are indicated by black downward triangles.

Statistically significant differences (Wilcoxon test, p *<*0.05) for syntax order *k* are indicated by solid downward triangles above the abscissa. ER values of continuous microstate sequences are lower than first-order syntax for word length *k* = 2 − 6 and the difference grows with increasing syntax order *k*. Analogous results are observed for microstate jump sequences. SE values of continuous sequences are larger than their first-order Markov level for *k* = 2 − 6. For jump sequences, SE decreases with syntax order *k* and is lower than the corresponding first-order syntax.

We next compared ER/SE syntax properties of EEG microstate sequences from n_AD_=25 subjects with AD diagnosis (Figure 4). Qualitatively, the findings were similar to the CN group. The AD group showed lower than first-order ER values for continuous (*k* = 2 − 6) and jump (*k* = 3 − 6) sequences, indicating additional syntactic structure. SE was higher than first-order syntax for continuous sequences (*k* = 2 − 6) and lower than first-order syntax for jump sequences (*k* = 2 − 6). For jump sequences, ER and SE coefficients at *k* = 1 were minimally larger than for first-order syntax.

**Fig. 4.**
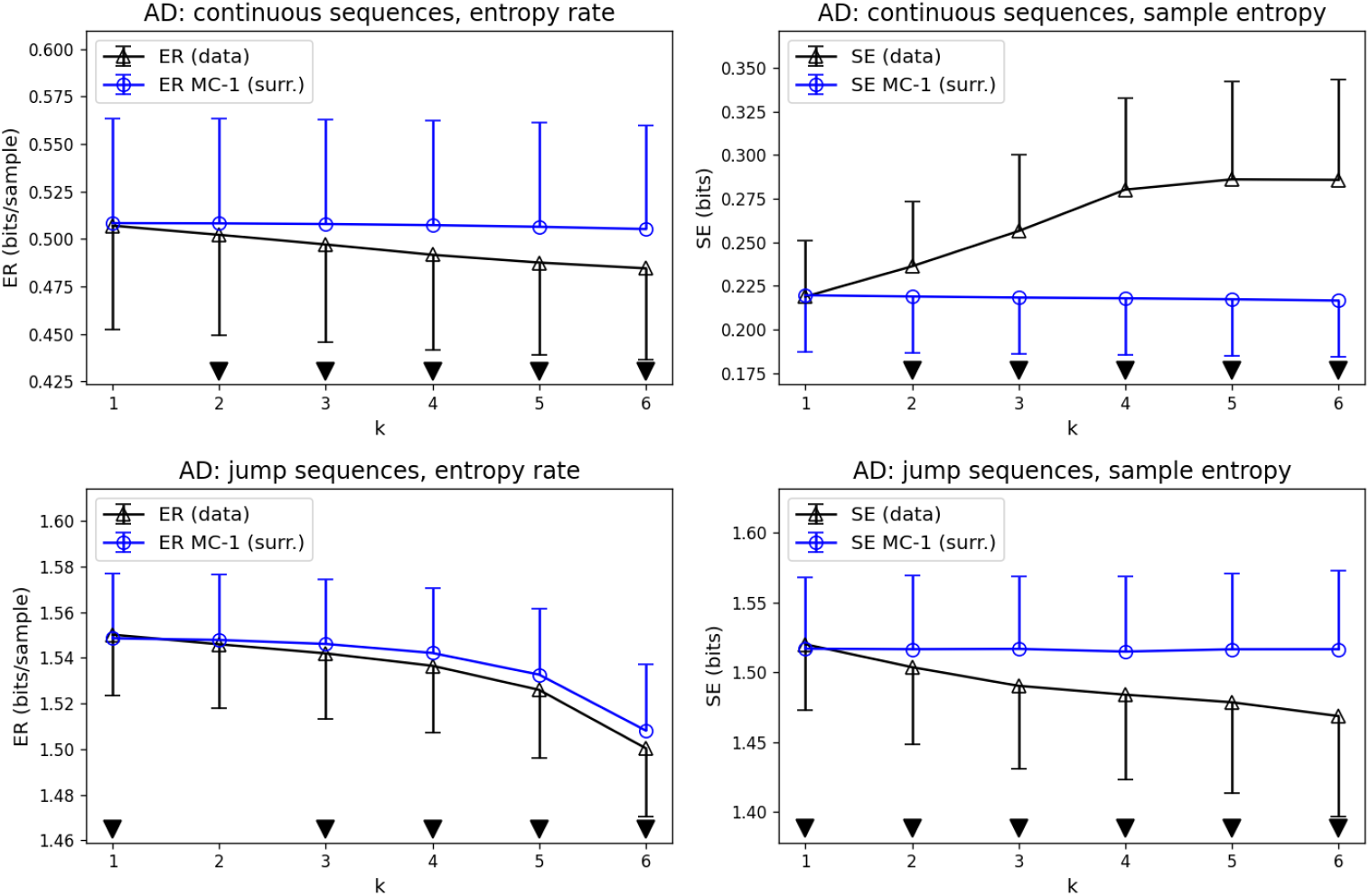
Entropy rate (ER) and sample entropy (SE) compared for real microstate sequences from the AD group (black, label: data) and their first-order Markov surrogates (blue, label: surr.). Statistically significant differences (p *<*0.05) between real microstate sequences and first-order Markov surrogates are indicated by black downward triangles.

The direct comparison of CN and AD subjects is shown in Figure 5. We found statistically significant results (p *<*0.05, Mann-Whitney U-test) for ER and SE at all syntax orders *k* = 1−6, for continuous and jump sequences. For continuous sequences, ER and SE were lower in the AD group. For jump sequences, ER and SE values were higher in the AD group. Visually, ER and SE curves had similar slopes and shapes in the CN and AD group, with the exception of SE in jump sequences, where the entropy curve appeared visually flatter in the AD group compared to CN. These observations will be analyzed quantitatively in the next section.

**Fig. 5.**
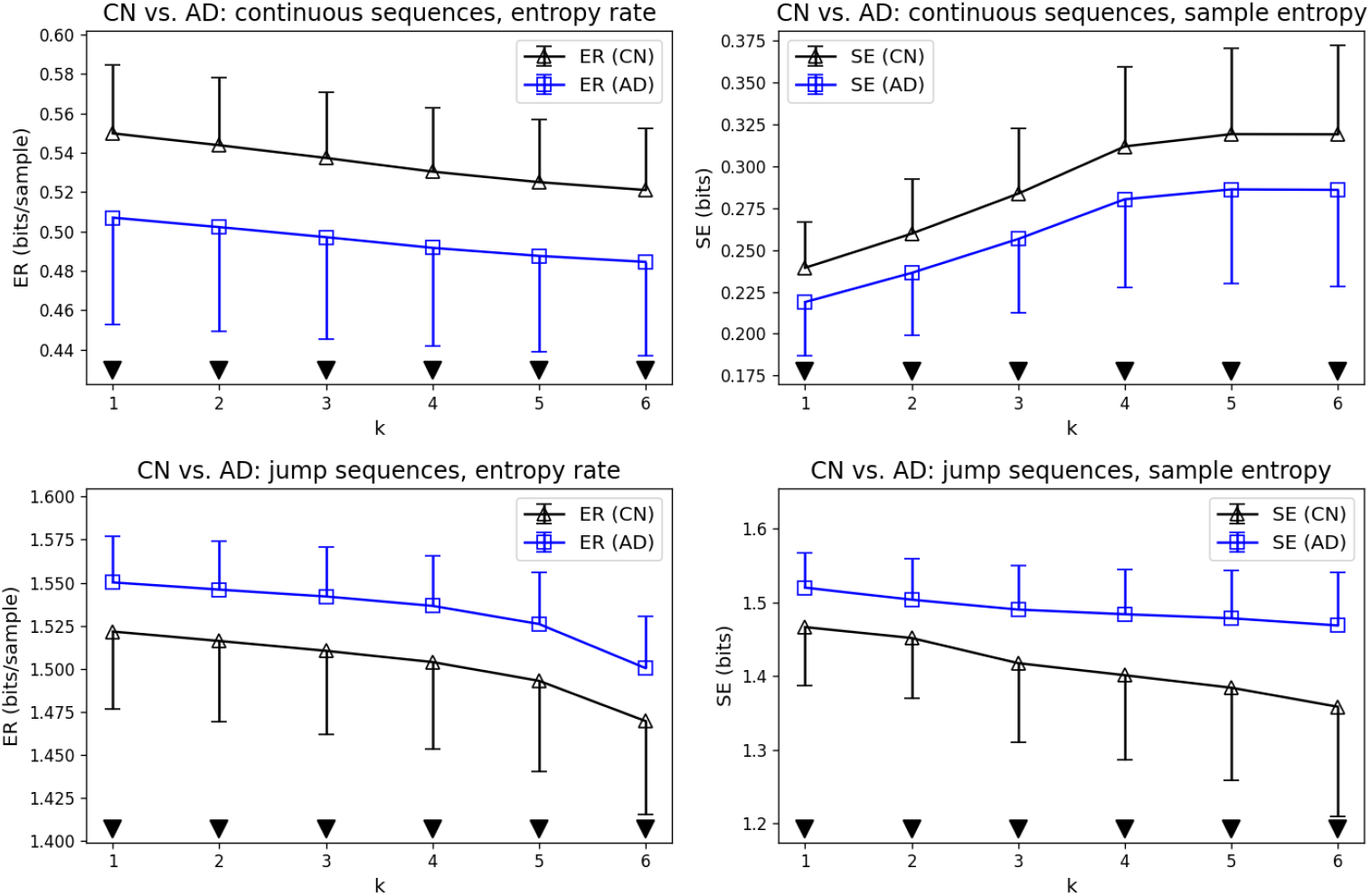
Direct comparison of CN and AD subjects. Statistically significant differences (p *<*0.05, black downward triangles) were found for all tested syntax levels (*k* = 1 − 6), for continuous and jump microstate sequences, and for both syntax metrics (ER and SE).

### 4.3 Normalized syntax

In Figure 5, it is observed that ER and SE curves were significantly different but had similar shape for CN and AD groups. To assess this shape independent of the absolute entropy values, we analyzed the normalized syntax measures defined for continuous (*σ*_*C*_) and jump (*σ*_*J*_) sequences defined above. It should be noted that larger values of *σ*_*C,J*_ indicate more syntactic structure, whereas large ER or SE indicate less syntax and more randomness. CN and AD group comparisons are shown in Figure 6. We found no statistically significant differences (p *<*0.05, Mann-Whitney U-test) for the normalized ER or SE syntax metrics between CN and AD groups at any syntax order *k* = 1 − 6.

**Fig. 6.**
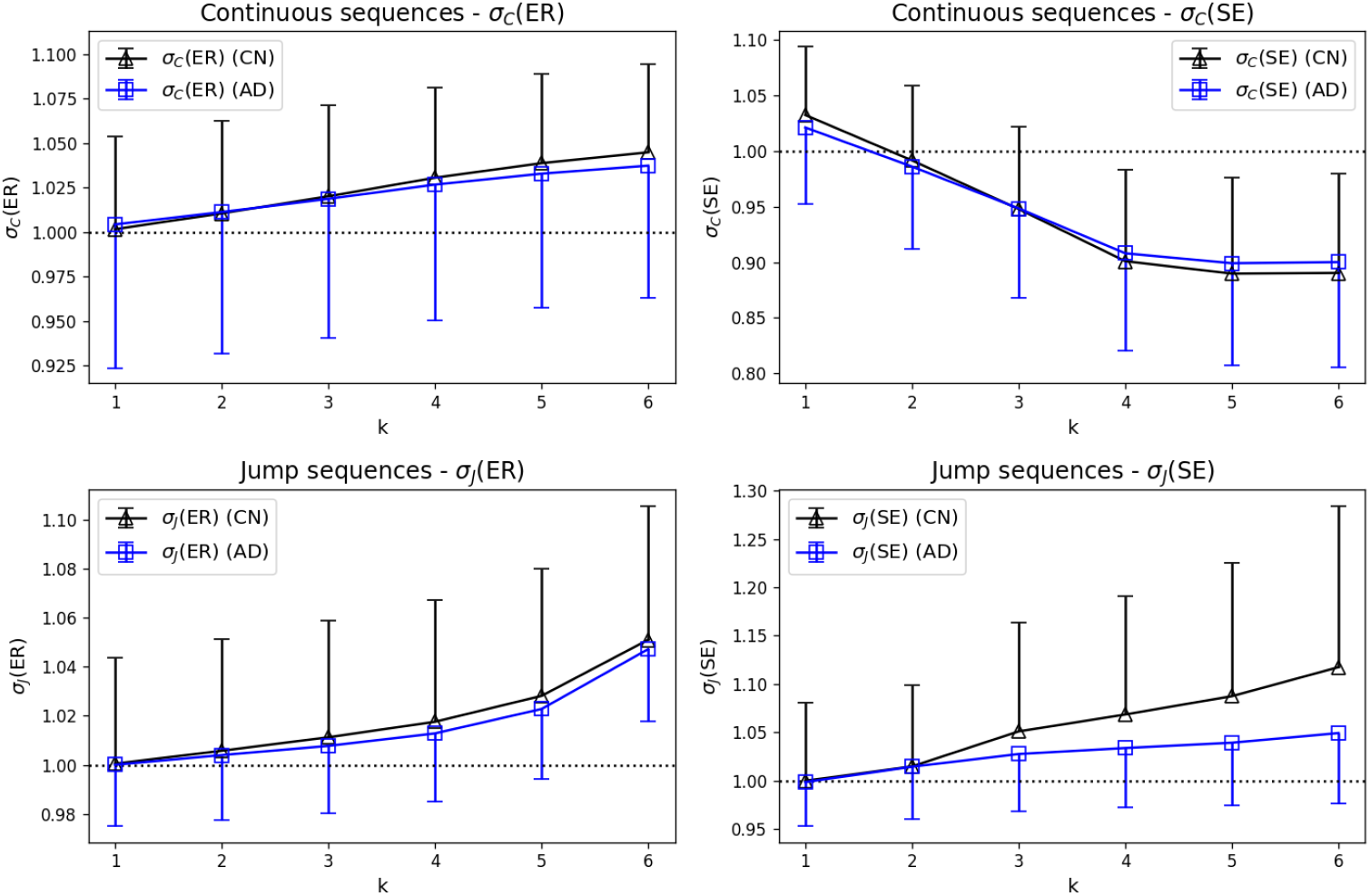
Normalized syntax measures *σ*_*C*_ and *σ*_*J*_ applied to ER (left) and SE (right) for CN (black) and AD (blue) groups. The first-order Markov syntax level is indicated by dashed lines (*σ*_*C,J*_ = 1). No statistically significant differences were found between CN and AD groups in continuous or jump microstate sequences.

## 5 Discussion

This study was inspired by a growing interest in the higher-order syntax properties of EEG microstate sequences (Haydock et al, 2025; von Wegner et al, 2025). The principal aim was to gain a better quantitative understanding of two different metrics that can be found in the literature, namely entropy rate and sample entropy (von Wegner et al, 2016, 2017; Wiemers et al, 2023; Hermann et al, 2024; von Wegner et al, 2023; Murphy et al, 2020; Artoni et al, 2022, 2023). We have given a detailed quantitative comparison of ER and SE and applied them to an EEG data set of individuals diagnosed with AD and controls. The main results are:

1. Theoretical ER and SE approximations for first-order Markov surrogates are closely related and yield an inequality ER (*k*) ≥ SE (*k*) for fixed microstate word size *k*. The inequality also holds for the empirical microstate sequences with non-Markovian properties investigated in this study.
2. Control and Alzheimer’s disease groups both show non-Markovian higher-order syntax properties.
3. Continuous and jump microstate sequences show different higher-order syntax when assessed by either ER or SE.
4. Higher-order syntax differs between control and Alzheimer’s disease groups.
5. Normalized ER and SE metrics show that certain higher-order syntax patterns are preserved in Alzheimer’s disease patients.

### 5.1 Theoretical ER and SE values for first-order syntax

Both metrics, ER and SE, have been applied to continuous dynamical systems in the past (Eckmann and Ruelle, 1985; Gaspard and Wang, 1993; Richman and Moorman, 2000; Lake et al, 2002). The fact that microstate sequences are discrete-valued sequences over a small alphabet allowed us to derive some quantitative results and to simplify the presentation and interpretation. For continuous systems, numerical estimates of these entropy variants contain additional parameters to measure the proximity of sample paths, and the theoretically exact entropies are defined as limits when these parameters approach either zero (proximity radius) or infinity (time steps). SE computations for continuous systems record a collision when 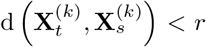, where d is a distance metric of choice, and *r >* 0 is a free parameter that defines the size of the collision volume. Entropy estimates are then reported for a range of parameter values and their consistency across the parameter range is tested (Gaspard and Wang, 1993; Richman and Moorman, 2000). The parameter *r* is not needed for discrete microstate sequences because the equality 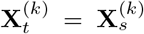 can be measured exactly, but some microstate studies do report this parameter.

Discrete variables also facilitate the analysis of Markov chain surrogates which are the null hypothesis against which higher-order syntax is tested (von Wegner et al, 2025). When first-order properties are assumed in the simplified expressions for ER (*k*) and SE (*k*), we obtain the theoretical approximations in Equation 5 and Equation 6. These approximations are useful for several reasons.

First, they provide a quantitative framework for ER and SE interpretation. Wu et al (2025) and Teng et al (2024) compare SE values between groups but in the absence of a ‘within group’ reference value, it is not clear whether the syntax structure of the microstate sequences in each group is richer than first-order or not. Murphy et al (2020) used shuffled microstate sequences as a baseline and show that jump sequences from control subjects have lower SE (more syntax) than shuffled surrogates. It can be shown that shuffled sequences have a first-order Markov structure, but it can also be demonstrated that their first-order properties are different from the original sequence, which complicates the interpretation of the results in terms of higher-order syntax (von Wegner et al, 2025).

Secondly, the theoretical approximations show that zero- and first-order syntax are reflected by flat ER and SE curves for *k* ≥ 1. This allows a concise reinterpretation of the results reported for individuals with psychosis in Murphy et al (2020). Their flat SE curve for *k* ≥ 1 suggests that these microstate sequences had a simple first-order Markov structure. This hypothesis could be tested independently with direct Markovianity tests (von Wegner et al, 2017; von Wegner and Laufs, 2018). In Murphy et al (2020) and Wu et al (2025), it can be observed that the first SE coefficient is significantly larger than subsequent SE coefficients. In both articles, the first coefficient has index 1. Following the classical SE definition (Richman and Moorman, 2000), this coefficient compares the recurrence of templates of lengths zero and one, i.e. *k* = 0. As there are no templates of length zero, algorithms (e.g. (Lake et al, 2002)) divide the number of single microstate recurrences (*X*_*t*_ = *X*_*s*_ for *t* ≠ *s*) by the total number of sample comparisons, i.e. *n* (*n* − 1) */*2 for a sequence of length *n*. We omitted this coefficient as our syntax concept starts with microstate word size *k* = 1. This explains why our SE curves are flat, as opposed to those in Murphy et al (2020) and Wu et al (2025). If needed, a good estimate of the SE coefficient for *k* = 0 can be obtained from the microstate distribution **p** alone and a temporal independence assumption (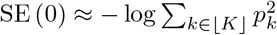, the second order Rényi entropy of **p**).

The third advantage of having a specific numerical value for the expected ER and SE values is the assessment of numerical inaccuracies due to limited sample size as illustrated for jump sequence ER estimates in Figure 9.

Lastly, we found a practical inequality ER (*k*) ≥ SE (*k*) for the Markov null hypothesis which also held true for the empirical microstate sequences investigated in this study. This inequality, in conjunction with the equivalence between Lempel-Ziv complexity (LZC) and ER reported in von Wegner et al (2023) (LZC ∼ ER ≥ SE), should facilitate the comparison of microstate studies combining these methods, such as the recent study by Teng et al (2024) who combined LZC and SE.

In summary, we strongly suggest that future studies employing ER or SE should either report the theoretically expected first-order syntax levels, or provide numerical estimates derived from first-order surrogates. If surrogate data is used, it should be synthesized from the actual transition matrix and not from shuffled sequences (von Wegner et al, 2025).

### 5.2 Sample size

Higher-order syntax metrics based on entropy can be data-hungry as they involve the estimation of high-dimensional discrete distributions (histograms) and the number of histogram bins grows exponentially with word length *k*. The effects are observed in Figure 9, where ER estimates from jump sequences display a bias towards lower values for *k* ≥ 4 that is corrected when the surrogate sequences are extended to the length of the underlying continuous microstate sequence. For real microstate sequences, the length is fixed and can become critically low for microstate jump sequences, i.e. sequences without adjacent duplicate labels. In this study, we used continuous sequences with 10^5^ samples and the resulting jump sequences were more than ten-fold shorter. Murphy et al (2020) processed three minutes of resting–state EEG sampled at 1 kHz and obtained jump sequences with approximately 2000-2500 samples, and Tait et al (2020) truncated jump sequences to a fixed length of 250 samples for complexity analysis. Some authors segment the EEG into 2 second epochs (Nishida et al, 2013; Grieder et al, 2016). At very short sequence lengths, a significant bias must be expected for higher-order syntax metrics. Likewise, we have observed that short sequences can cause theoretically flat SE curves to increase slowly with *k*, as seen in Wu et al (2025). To account for these effects, we suggest again the use of Markov surrogates to distinguish real higher-order effects from biases due to sample size.

### 5.3 The meaning of ‘the entropy of a microstate sequence’

Our results show that there is no single answer to the question ‘what is the entropy of a microstate sequence?’, without providing further methodological details. Studies using a single entropy metric, e.g. ER or SE, often discuss their results just in terms of ‘entropy’, and this can be easily misunderstood as a unique entropy concept to which different estimators are applied. Although some metrics coincide, ER and SE refer to different theoretical concepts and, as we have shown, ER and SE do not necessarily increase or decrease in parallel. We have presented a similar situation in von Wegner et al (2023), where we chose different types of entropy to distinguish Kolmogorov complexity from statistical complexity.

Review articles on complexity analyses for schizophrenia Fernández et al (2013) and across neuro-psychiatric diseases, including Alzheimer’s disease Takahashi (2013), show that contradictory findings are not uncommon when different entropy measures are employed. One should therefore not expect a simple answer to the question whether a neurodegnerative condition like AD causes a decrease or an increase in syntactic complexity.

Figure 3 and Figure 4 in this manuscript illustrate the importance of two factors, the type of microstate sequence (continuous vs. jump) on the one hand, and the entropy metric (ER vs. SE) on the other. In our dataset, continuous microstate sequence syntax is classified differently by ER and SE. ER coefficients are below the first-order Markov level and thus indicate more syntactic structure than encoded in the transition matrix, whereas SE values lie above the first-order syntax level. We explain these opposite outcomes by the different algorithmic approaches. ER (*k*) considers a relatively short time window into the past of the current microstate, whereas SE (*k*) scans the entire sequence for recurrences of the microstate word 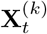. ER is therefore dominated by the sequence’s short-range autocorrelations and these contribute constructively to short-term predictability and effectively lower the (finite) entropy rate. SE, on the other hand, can be affected by non-stationarities, especially when long time series as used in this study (10^5^ samples, 200 seconds) are considered. Microstates recurring with a slightly different duration, for example, will effectively increase SE coefficients. Other non-stationary features in longer EEG recordings can be due to drowsiness or varying cognitive engagement during the assumed resting state. Low SE values indicate a higher chance of finding exactly the same microstate word at different time points. We would not call this effect self-similarity as in Murphy et al (2020), however, as this would indicate the recurrence of similar microstate patterns at different time scales, not different positions on the time axis (Embrechts, 2009). Re-scaling, i.e. dilation or compression of a time series, which is at the core of self-similarity, is well defined for real-valued time series, but not for nominal variables such as microstates.

The second factor that determined whether we found more or less syntactic structure was the type of microstate sequence. Jump sequences showed ER and SE coefficients below the first-order syntax level, i.e. they had more syntax than expected from the transition matrix. This demonstrates that there is a predictable higher-order structure in the sequence of brain states when the duration of the individual state is ignored.

Taken together, these results illustrate the importance of precisely documenting the pre-processing and analysis pipeline applied to EEG data. Unfortunately, the microstate literature is not always explicit about the sequence type (continuous or jump) that was analyzed. Statements like ‘microstate sequences were obtained by back-fitting the template maps into the EEG dataset’ are not precise enough to reproduce results and compare syntax changes across studies.

### 5.4 Microstate syntax and EEG complexity in Alzheimer’s disease

Several studies have analyzed microstate dynamics in AD but not all findings have been concordant. Different studies have found microstate duration to be shorter (Strik et al, 1997; Dierks et al, 1997), unaltered (Nishida et al, 2013; Musaeus et al, 2019), or longer (Smailovic et al, 2019; Tait et al, 2020; Musaeus et al, 2020). These differences have been attributed to microstate processing parameters and to patient characteristics such as AD stage and cholinergic medication (Musaeus et al, 2020; Lian et al, 2021). Longer microstate duration is possibly related to more advanced disease stage and medication is assumed to normalize certain EEG parameters (Lian et al, 2021).

First-order microstate syntax analysis using the method proposed by Lehmann et al (2005) did not reveal any differences in AD patient groups in several studies by (Nishida et al, 2013; Musaeus et al, 2019; Tait et al, 2020), whereas altered transition probabilities for individual microstates (canonical microstates A, B) were reported in other studies (Musaeus et al, 2020; Lian et al, 2021). Negative first-order syntax analysis results motivated Tait et al (2020) to apply Lempel-Ziv complexity (LZC) which revealed decreased complexity of microstate jump sequences from the AD group. As reported in von Wegner et al (2023), LZC and ER measure the same complexity/syntax property and converge towards the same numerical value for long sequences. Our results therefore contradict those in Tait et al (2020), as we found increased ER for jump sequences in the AD group. This difference is not explained by the algorithmic approach as LZC analysis of our data gave the same qualitative result (data not shown), i.e. more randomness and less syntactic structure in AD jump sequences. A major difference between the studies is that Tait et al (2020) analyzed 20 seconds of eyes-open EEG and truncated all jump sequences to 250 samples, whereas we analyzed 200 seconds of eyes-closed EEG resulting in average jump sequence lengths of 8564 samples (CN) and 7667 samples (AD). Following the argument given further above, we believe that the extended observation time allows all three algorithms (ER, LZC, SE) to detect drifts and fluctuations in brain state sequences that occur more randomly in AD. The differences between eyes-open and eyes-closed conditions might also have contributed to the different outcomes. In agreement with our findings, other complexity measures computed from broadband and multichannel EEG, and in that sense similar to the microstate approach, found increased complexity in AD patients (0.3-60 Hz (Yoshimura et al, 2004) and 0.5-25 Hz (Czigler et al, 2008)). Another recent study compared short microstate sequences (2 second epoch length) between individuals with mild cognitive impairment (MCI) and controls, and reported no significant differences in SE coefficients (*k* = 1 − 7) (Wu et al, 2025). Nevertheless, they achieved successful classification combining basic microstate parameters, SE coefficients, and another entropy measure in a machine learning approach (Wu et al, 2025).

In summary, similar to basic microstate statistics, divergent results are not uncommon in EEG complexity analyses from different AD cohorts, as reviewed in Takahashi (2013).

Another novel finding of our study is the distinction between absolute entropy differences and the shape of the entropy (ER/SE) curve when plotted against the syntax order *k*. ER and SE differences between control and AD groups (Figure 5) appeared as a near-constant ‘offset’ between CN and AD, whereas the shape of ER and SE curves was largely similar. The slope of the ER/SE curves, as a measure of the increasing difference between first-order and higher-order properties might itself prove to be a useful tool to quantify brain state dynamics in the future. Our current analysis with the newly introduced syntax measure *σ* (*σ*_*C*_ for continuous and *σ*_*J*_ for jump sequences) suggests that brain state dynamics in the AD group retain higher-order syntax patterns to some degree, while evolving at a higher absolute randomness level. While our normalized syntax metric did not reveal statistically signficant differences between AD and control, the visual appearance of the curves suggests a hypothesis that AD syntax curves eventually might turn out to be flatter if analyzed on a larger data set.

## 6. Conclusion

Our results show that continuous and jump microstate sequences from control and AD individuals display higher-order syntax structure, making these features a possibly valuable biomarker of brain health. From a methodological perspective, our results emphasize the need for long input sequences and low-order Markov surrogates to reliably estimate higher-order syntax properties and possible sources of bias.

## Declarations

This work was funded by the German Research Foundation (DFG), project number 440536202.

**Fig. A1.**
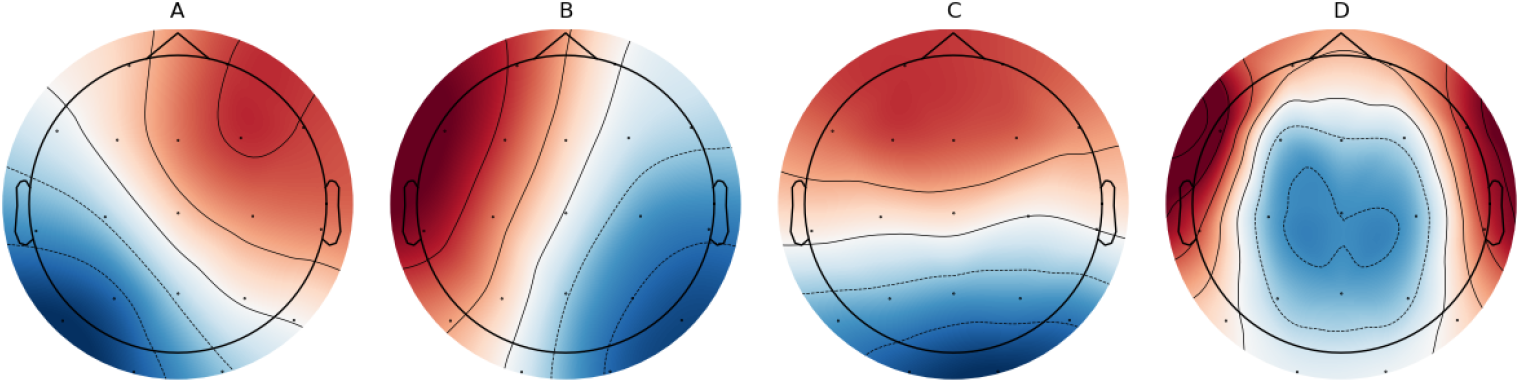
Group-level microstate maps from the modified K-means algorithm. Subject-level maps were obtained from eyes-closed resting-state EEG recordings of control subjects and individuals with Alzheimer’s dementia, as published in Miltiadous et al (2023).

## A Appendix

### A.1 Microstate analysis

This section presents the basic microstate statistics and the group-level microstate maps for K=4 are shown in Figure A1. The microstate statistics duration, occurrence, and coverage are given in Table A1. It should be noted that the qualitative results in the main text are not dependent on the number of clusters *K*, or the exact numerical outcomes of the microstate clustering and fitting algorithm.

**Table A1.** Microstate statistics duration, occurrence, and coverage for microstate (MS) classes A-D presented as means (standard deviation) for the CN and AD groups. The p value of the Kruskal-Wallis test between the CN and AD groups is shown in the fourth column, values p *<*0.05 were considered statistically significant.

| MS | CN | AD | p value |
| --- | --- | --- | --- |
| Duration (ms) |  |  |  |
| A | 22.70 (2.03) | 25.66 (4.89) | **0.024** |
| B | 19.69 (1.47) | 26.38 (6.50) | **<0.001** |
| C | 28.69 (4.33) | 30.10 (6.00) | 0.631 |
| D | 19.63 (1.72) | 23.96 (3.60) | **<0.001** |
| Occurrence (1/s) |  |  |  |
| A | 11.57 (1.03) | 9.48 (1.47) | **<0.001** |
| B | 9.29 (1.34) | 9.34 (1.54) | 0.810 |
| C | 13.35 (0.91) | 10.82 (2.10) | **<0.001** |
| D | 8.54 (1.60) | 8.38 (2.25) | 0.562 |
| Coverage (%/100) |  |  |  |
| A | 0.26 (0.04) | 0.24 (0.03) | **0.036** |
| B | 0.18 (0.04) | 0.24 (0.06) | **0.001** |
| C | 0.38 (0.07) | 0.32 (0.07) | **0.002** |
| D | 0.17 (0.04) | 0.20 (0.05) | 0.093 |

